# Pep2Vec: An Interpretable Model for Peptide-MHC Presentation Prediction and Contaminant Identification in Ligandome Datasets

**DOI:** 10.1101/2024.10.14.618255

**Authors:** William John Thrift, Quade Broadwell, Jason Perera, Nicolas W. Lounsbury, Jieming Chen, Suchit Jhunjhunwala

## Abstract

As personalized cancer vaccines advance, precise modeling of antigen presentation by MHC class I and II is crucial. High-quality training data is essential for clinical models. Existing deep learning models focus on prediction performance but lack interpretability. We introduce Pep2Vec, a modular, transformer-based model trained on MHC I and II ligandome data, transforming input sequences into interpretable vectors. This approach integrates source protein features and elucidates the source of its performance gains, revealing regions that correlate with gene expression and protein-protein interactions. Pep2Vec’s peptide latent space shows relationships between peptides of varying MHC class, allotype, lengths, and submotifs. This enables identifying four major contaminant types, constituting 5.0% of our data. Pep2Vec enhances MHC presentation prediction, achieving higher average precision on our presentation test set and immunogenicity datasets than existing models, and reducing contaminant-like peptide recommendations. Pep2Vec addresses a critical need for the development of more precise and effective applications of peptide MHC models, such as for cancer vaccines and antibody deimmunization.

## Introduction

Translating deep learning models into clinical applications requires a focus on interpretability to ensure trust and efficacy. FDA has emphasized the importance of trustworthy AI for drug discovery by stating that "an AI/ML system may exhibit limited explainability due to its underlying complexity… These concerns have resulted in a focus on developing standards for trustworthy AI that address specific characteristics in areas such as explainability…”^1^ With the translation of deep learning models for the selection of cancer vaccines to the clinical setting,^2,3^ this field has the opportunity to blaze the trail for interpretable drug discovery models.

The adaptive immune system’s ability to target cancer relies heavily on the presentation of tumor-derived peptides by MHC molecules. MHC class I (MHCI) molecules present peptides to CD8+ T cells, critical for the direct elimination of tumor cells, while MHC class II (MHCII) molecules activate CD4+ T cells, essential for modulating the immune response.^4^ These processes are fundamental to the efficacy of cancer vaccines and personalized therapies such as individualized neoantigen specific therapies (iNeST).^5–7^ However, accurately predicting which peptides will be presented by MHC molecules is challenging due to experimental constraints that can lead to the detection of peptides that are not truly presented.^8^ Sophisticated deep learning models trained on such spurious peptides have sufficient capacity to learn from these examples, and may give them favorable predictions.^9^ This behavior is a major challenge for the clinical use of peptide-MHC (pMHC) presentation models, where valuable slots in personalized cancer vaccines should not be wasted on non-presented peptides. Interpretable models can enable the identification of these experimentally observed but non-presented peptides and to enhance our understanding of pMHC presentation, analogous to such work in other fields.^10–12^ Achieving these goals is crucial for developing more effective cancer vaccines, and deimmunizing antibody drugs, among other pMHC presentation applications.

While biophysical properties of MHC ligands primarily influence their binding to MHC, there are other biochemical factors that contribute to quantitative modulation of pMHC presentation, like peptide processing and gene expression. Efforts to extend pMHC models to account for these and other factors have evolved significantly. Early efforts included modeling proteasome cleavage by incorporating residues from the source protein flanking the cognate peptide.^13–15^ Incorporation of gene expression as a feature has also been explored^16–18^. Furthermore, our earlier work on integrating source protein features derived from protein language models (PLMs) has enhanced MHC class I (MHCI) predictions by capturing a broader range of determinants, notably and somewhat surprisingly serving as proxies for gene expression.^19^ However, these studies have not employed the comprehensive end-to-end training that has proven effective in deep learning, nor have they elucidated how these protein features contribute to improved model performance, leaving their limitations and full potential still unexplored.

Ligandome data for training pMHC presentation models are typically generated using mass spectrometry to identify ligands that are eluted from immunoprecipitated pMHC complexes. This method has some well known biases and sources of noise as well as a number of less well known or one-off sources of error that can confound downstream analysis.^20,21^ Some well-known biases include underrepresentation of cysteine and existence of common contaminant proteins (such as the “crapome”^22^). Besides biases related to the assay, other issues may affect large-scale ligandome data. For example, misannotations of MHC allotypes in some samples, incomplete abrogation of endogenous MHC genes in engineered cell lines used for single-allele data generations, etc. Finally, there are a host of additional factors that may be related to either the individual experiment, search, or filtering, that can introduce error.^23,24^

An interpretable peptide space can systematically reveal contaminant peptides or QC issues by leveraging patterns within this space. Previously, Sarkisova et al.^16^ developed a method to create peptide latent spaces that could identify submotifs; however, this approach was limited to a single MHC allele and peptide length at a time. This limitation hindered its ability to analyze trans-MHC peptide relationships that could help identify contaminants, and the approach was never extended to MHCII. In contrast, while Gibbs Clustering,^25^ is applicable across different peptide lengths and MHC classes, it lacks the structured metric space offered by the Sarkisova method. Recently, MHCpLogics^26^ created a multidimensional scaling based - tool (similar to the Sarkisova method) for identifying submotifs which overcomes the limitation of single MHC alleles and single peptide lengths, but the application has been limited to a maximum of 6 alleles and only MHCI.

In this work, we introduce Pep2Vec, a modular, transformer-based model specifically engineered to create interpretable vectors for each input sequence. The model’s peptide submodule generates interpretable vectors and locates binding cores using specialized attention mechanisms. Additionally, our approach mitigates the risk of overfitting across diverse MHC alleles by employing dimensional reduction and protein language models on the MHC amino acid sequences, ensuring a comprehensive representation of MHC variability. The model’s source protein submodule significantly enhances performance, particularly for MHCII, by integrating source protein features that enable a deeper understanding of pMHC presentation, leading to improvements on our presentation test set compared to existing models, as well as several immunogenicity datasets. This integration is evidenced by precise mappings and analyses within the Source Protein Vector space, which reveal distinct clusters associated with gene ontologies and local structure dictated by protein-protein interaction networks. Moreover, the Peptide Vector space reveals the relationship between peptides regardless of length, presenting MHC allele, and MHC class. It also elucidates relationships among peptides of various lengths, presented MHC alleles, and MHC types. This framework has enabled the systematic identification of four contaminant types—missanotation of a sample’s alleles, MHCI contaminants in MHCII ligandomes, novel protein-terminus peptides, and low-complexity peptides—that together account for 5.0% (77,475 peptides) of our ligandome data. Collectively, these advances establish a new benchmark in MHC presentation prediction and prevent the nomination of spurious peptides as therapeutic candidates or biomarkers in immunotherapies or biomarkers in immunotherapies, and deimmunization of large molecule drugs.

## Results

### Pep2Vec Model Architecture and Performance

Pep2Vec is a modular, transformer-based model that is designed to generate interpretable vectors representing each input sequence, as depicted in Figure 1 a). The model incorporates two types of inputs: 1) Features related to the peptide (peptide sequence, source protein features, and flanking sequences). For each of these features, a vector representation is learned via self-attention and combined using an elementwise addition operator, forming the Integrated Peptide Vector. This operation ensures independence across the vector spaces of each feature type. 2) Features describing the MHC allele. A vector representation is also learned for the MHC allele and combined with the Integrated Peptide Vector using an elementwise multiply operation. This operation enforces correspondence between the MHC Vector and the Integrated Peptide Vector. The resulting vector is processed through a multilayer perceptron to predict the presentation likelihood of the peptide-MHC (pMHC) complex, as illustrated in Figure 1 a).

**Figure 1:**
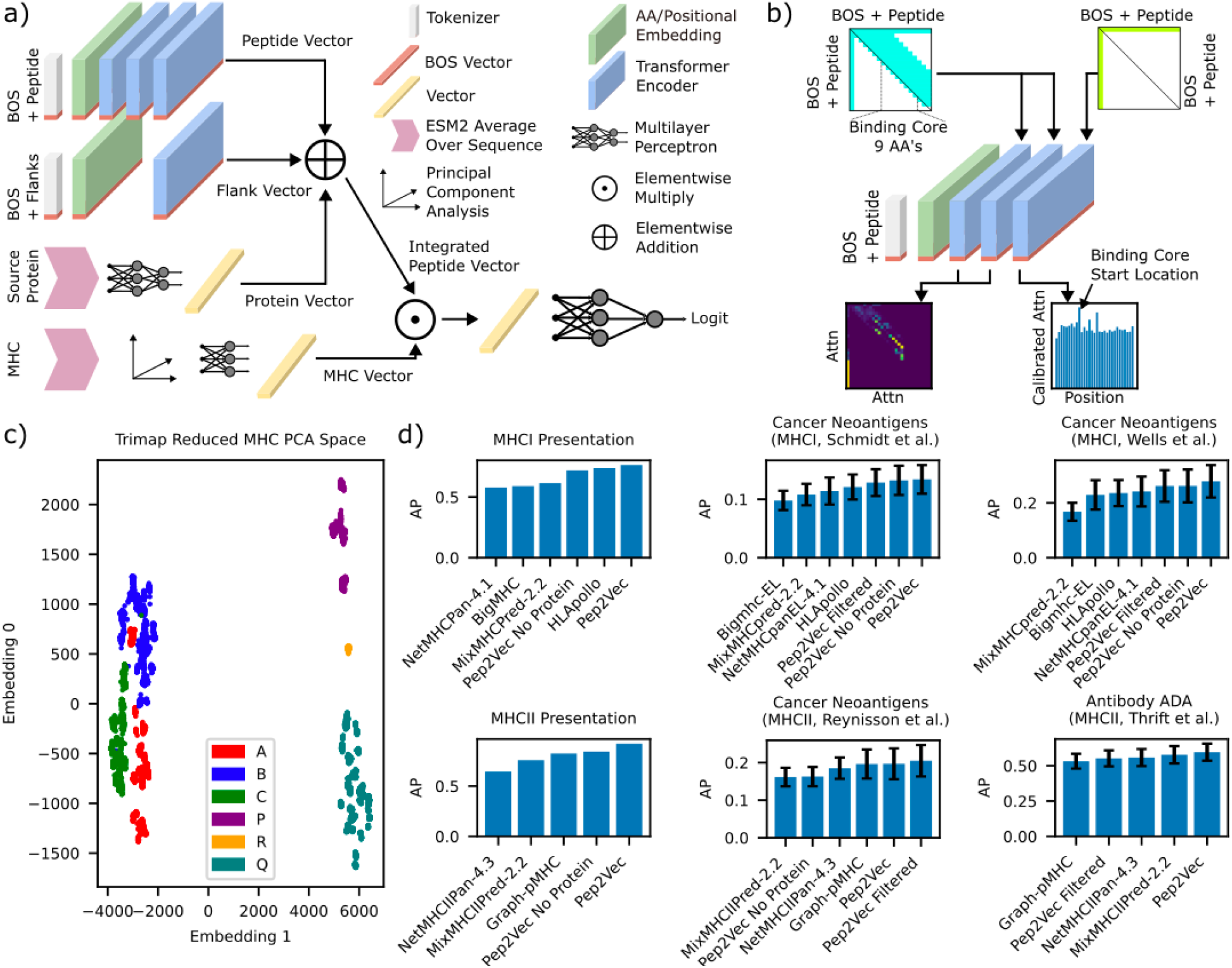
Pep2Vec model diagram and performance. a) A diagram of the deep neural network used through the work. BOS: Beginning of sequence, ESM: evolutionary scale model.

The peptide submodule (depicted in Figure 1 b)) serves two primary functions: 1) It generates an interpretable vector that represents the peptide. This is achieved by concatenating a Beginning of Sequence (BOS) token to the input peptide sequence. The vector associated with this token, which is the only information passed to the rest of the model from this submodule, is updated by transformer blocks to incorporate the rest of the peptide information. A specialized attention mask (shown in neon green in Figure 1b)) is applied to enforce this targeted behavior in the model. 2) It produces attention values from which the starting location of the binding core for MHC class II (MHCII) peptides can be derived. To achieve this, we introduce a binding core inductive bias into the model. This is done by modifying the attention masks of the transformer to only allow residues within 9 amino acids (the length of the MHCII binding core) of each other to update a particular residue’s information (allowed residues depicted in neon blue in Figure 1b)). This configuration of the peptide submodule creates cross-attention between the BOS token and the peptide residues at the end of the submodule. After calibration (see methods), the largest attention value is obtained at the binding core starting position, as shown in the attention histogram in Figure 1b.

Peptide Presentation by MHC is influenced by factors beyond the peptide sequence, such as source protein processing and expression. We go beyond earlier works by incorporating features that represent the entire source protein. This is achieved by obtaining protein language model (PLM) features (specifically ESM2) of the source protein and averaging these features residue-wise to obtain a single-vector representation. Useful features from these vectors are subsequently extracted using a multilayer perceptron (Figure 1 a)). This approach enables the model to capture features related to the source protein that indirectly modulate presentation of the peptides from a protein, such as protein compartmentalization and correlations between protein families and the frequency of peptide presentation. The inclusion of source protein information requires a test-train split with unseen proteins in test, following the split strategy developed by Graph-pMHC,^27^ but here we also modify our negative sample method to ensure no data leakage due to the source protein, see methods. Below, we explore these factors by interpreting the Source Protein Vector.

An important challenge of pMHC modeling is the relatively limited diversity of MHC alleles (data has been collected for just a few hundred out of thousands of possible alleles), which can potentially lead to overfitting for pan-allelic models which process the entirety of the MHC sequence, or even just a pseudosequence. Here, we greatly reduce the number of model parameters which are fit to represent the allele sequence through the use of PLMs and parameter-free dimensional reduction applied using the entire observed MHC space rather than just those for which pMHC data has been acquired. First, we generate an ESM2-derived vector representation for the MHC protein, following the procedure used for the source protein, to capture MHC variability. Next, we apply a principal component analysis (PCA) dimensional reduction to all 15,175 MHC alleles/allotypes that have been observed and uploaded to IMGT.^28^ This full MHC space, further reduced to 2D for the purpose of visualization by Trimap^29^, is shown in Figure 1c. By training using the observed MHC space, we ensure that this dimensional reduction can capture alleles outside of our training dataset. Finally, a single linear layer (accounting for all learned MHC parameters) maps the output vector into the MHC Vector.

Pep2Vec is jointly trained on MHCI and MHCII, which, while producing limited benefit for model performance (see Supplemental Figure 1 for a full ablation study of Pep2Vec), is useful for model interpretation as shown below. Because of this, we compare Pep2Vec’s performance with various recent MHCI and MHCII models. We find that Pep2Vec outperforms contemporary models on pMHC presentation datasets.^2,19,27,30–33^ Pep2Vec also achieves the best performance on a range of immunogenicity datasets: two MHCI neoantigen immunogenicity datasets, a MHCII neoantigen dataset, and a clinical Antibody anti-drug antibody (ADA) (MHCII immunogenicity driven) dataset. We found that protein features consistently improved performance on immunogenicity datasets for both MHCI and MHCII. A barchart depicting these performance comparisons is shown in Figure 1 d). We note that for clinical antibody ADA, MHCII presentation models outperform models that seek to evaluate tolerization alone, such as the recent AbNatiV,^34^ which reports a PPV at 80% sensitivity of 38%, while Pep2Vec achieves 56% PPV at this sensitivity.

Sequences representing the peptide and the residues flanking the peptide from the source protein are processed with transformer layers. A learned source protein representation is derived from the source protein’s ESM embedding. The peptide, flank, and source protein representations are added together to form the peptide representation. A learned representation of the MHC allele is derived from a PCA dimensionality reduction of the MHC allele’s ESM embedding. The peptide and allele representations are then compared via an elementwise multiplication yielding a vector representation of the full input, and a multilayer perceptron is used to obtain the input’s presentation likelihood. b) A more detailed schematic of the peptide processing transformer. Attention masking in the early transformer layers is used to create an inductive bias favoring the use of binding core level representations (shown in the lower left attention map). In the final transformer layer, attention masking is used to simplify the attention to just BOS - token interactions. This is shown in the lower left attention mask, where the token with the largest BOS - token attention is interpreted as the binding core starting location for MHCII peptides. c) A 2-D trimap dimensional reduction of the MHC PCA space (global score 84%), showing the relationship between different MHC. d) Barcharts depicting the performance (measured with average precision (AP)) on the MHCI presentation test set used here (# positives: 84,916), CD8+ cancer neoantigens from Schmidt et al.^35^ (# positives: 127) and Wells et al.,^36^ (# positives: 41) the MHCII presentation test set used here (# positives: 88,398), CD4+ cancer neoantigens from Reynisson et al.,^37^ (# positives: 55) and a clinical antibody anti-drug antibody immunogenicity, (# positives: 43) dataset from Thrift et al.^27^ Immunogenicity datasets are bootstrapped over 500 evaluations due to the small number of positives.

### Interpretation of the Source Protein Vector

Above we found that the use of source protein features contributed significantly to our performance. Supplemental Figure 2 a) depicts precision-recall curves for the flagship Pep2Vec model and a version trained without protein features, with particularly notable gains observed for MHCII. We briefly note that this performance gain is due to generalization on novel proteins as we ensure no protein overlap between test and train, see methods. In the following section, we use the Source Protein Vector to interpret what information Pep2Vec is using to obtain these gains.

We create a 2D map of the Source Protein Vector space using a Trimap dimension reduction, as shown in Figures 2a-e. For Figure 2a, each protein is colored based on the average logit score of all peptides, both positive and negative, from our test set for each protein. Due to fewer negative examples for each positive in MHCII compared to MHCI, MHCII has higher average scores. The map reveals two primary clusters, with scores generally increasing with increased embedding #0. Notably, the bottom cluster shows a preference for high scores for MHCII.

**Figure 2:**
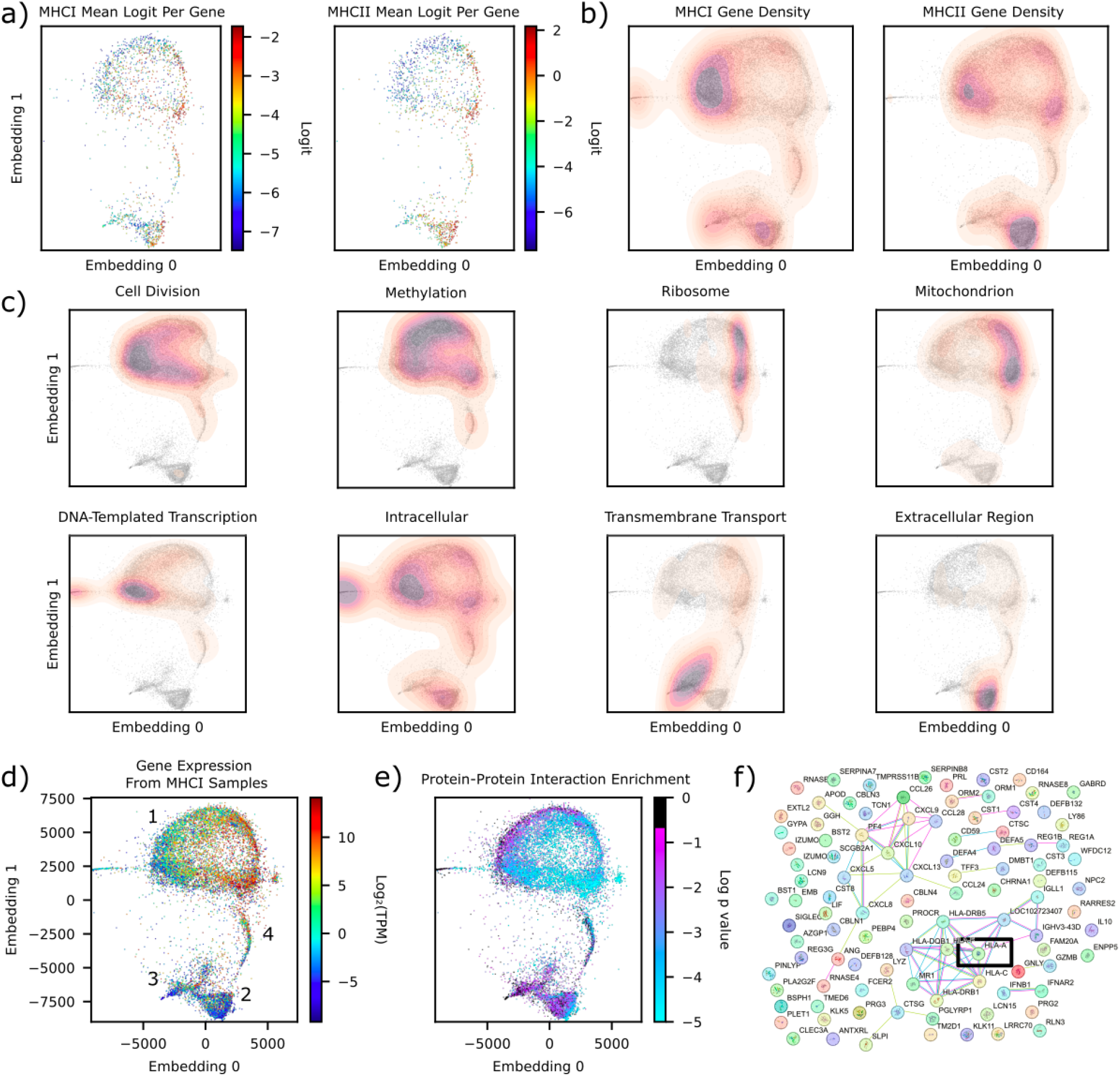
Pep2Vec’s learned source protein representation. a) A density plot of a trimap dimensional reduction of the source protein embeddings with only test set proteins (global score 79%). The color depicts the mean MHCI (left) and MHCII (right) logit score of all test set peptides for each protein, averaged by the proteins at each particular pixel of the plot. b) Contour maps of source protein density observed in the MHCI (left) and MHCII (lower) datasets The full source protein embedding is shown as a gray pixels for reference. c) Contour maps of source protein density observed for various gene ontologies. d) A density plot of a trimap dimensional reduction of the source protein embeddings with all proteins in our dataset. The color depicts the log transformed expression value of each protein, averaged by the proteins at each particular pixel of the plot. Expression values are themselves an average of the expression values observed across various MHCI datasets. e) A density plot of a trimap dimensional reduction of the source protein embeddings with all proteins in our dataset. The color depicts the log transformed p value of enriching for experimental protein-protein interactions (PPI) with the 100 nearest neighbors of the protein. f) A PPI graph obtained for HLA-A (dark square) for its neighborhood. Plot created using the stringdb tool subsetting to only experimental PPIs, in this plot, nodes are organized to highlight clusters, and not based on the position in the trimap embedding space.

Our investigation into the model’s organization of proteins within the Source Protein Vector space leads us to an intriguing observation. We first examine the enrichment of different protein types in various regions of the latent space. We present the density of proteins associated with MHCI and MHCII peptides in Figure 2 b). One may observe that MHCII peptides are primarily found from proteins in the bottom Cluster 2 (see Figure 2 d)), while MHCI peptides are more associated with the top Cluster 1. The long Cluster 4 between the MHCI and MHCII clusters exhibits roughly even proportions of peptides presented by both.

Next, we investigate the enrichment of proteins with various gene ontologies. We find that the primary MHCII cluster is, unsurprisingly, associated with peptides derived from extracellular proteins, while the small cluster just to the left of it is associated with transmembrane transport proteins. Cluster 3, just to the left of Cluster 2 on the left is associated with intracellular proteins. For the large MHCI dominant cluster, the right hand side is dominated by the mitochondrion (lower), and the ribosome (upper), which receive high mean logits, while the left hand side is dominated by proteins associated with cell division, which receive low mean logits.

The organization of the MHCI cluster is clarified by Figure 2 d), which shows the gene expression of each protein averaged over MHCI ligandome samples (which collected gene expression). A striking gradient of gene expression is apparent across the upper primary cluster, mirroring the logit scores depicted in Figure 2 a). This pattern is particularly noteworthy as gene expression is regulated by various promoters and other regulatory mechanisms, which are not encoded in the amino acid sequence of the source protein, but (based on Figure 2 c)) are correlated with the gene ontologies of the protein. We note that such organization by gene expression is not observed in the original ESM latent space, see Supplementary Figure 3, implying that this is learned only in the context of pMHC presentation While globally the organization of proteins in the Source Protein Vector space is dictated by their typical gene expression and presenting MHC, we find impressive local structure as well. Figure 2 e) depicts the Protein latent space colored by the statistical significance of local protein-protein interactions (PPI). Briefly, each protein in the Protein latent space is taken with its 100 nearest neighbors and the statistical significance of the enrichment of experimental PPIs among these 101 proteins from stringdb^38^ is determined, compared to the overall space of PPIs in stringdb. From this, one may observe that locally proteins are ordered by the other proteins they are associated with. We examine one neighborhood, that of HLA-A, in Figure 2 f). One may observe in this neighborhood that HLA-A has 8 experimental PPIs, with many genes related to the immune response being found in this neighborhood. We observe that local clustering is not influenced by protein structure, which might otherwise explain the colocalization of gene ontologies (e.g., membrane proteins sharing structural components easily captured by PLMs). To support this, we calculated the TM-align mean evalue for a protein with its 100 protein neighborhood. Selecting 1000 random proteins and calculating the mean evalue in this way, we find an evalue of 3.2 +/- 1.1, indicating chance structural similarities.

### Interpretation of the Peptide Vector

Pep2Vec’s Peptide Vector space condenses input vectors into presentation-oriented representations that highlight the holistic relationship between submotifs while disregarding extraneous information like peptide length and non-anchor residues. The complete Peptide Vector space is displayed in Figure 3 a), with specific zoom-ins for MHCI (Figure 3b)) and MHCII (Figure 3c)). Distinct clusters can be observed (particularly distinct for MHCI), which are dictated by the presenting allele. For MHCI, we observe a correlation of 0.37 between a distogram of the centroid locations of an allele’s peptides in the peptide space, and the distogram of the alleles in an allele space that uses blosum embedding of their psuedosequences. A closer look at a particular allele, for example, A*02:01 (Figure 3 d)), reveals the space’s fine structure of submotifs. Similar peptides are grouped closely together, and interestingly, these submotifs are clustered based on anchor residues, not length. This observation is demonstrated by the boxplot of the length composition for each cluster, which shows that most cluster’s length compositions closely follow that of the overall data (Figure 3 e)). Notably however, there are some clusters which deviate strongly from this, with one cluster being exclusively occupied by 8mers, and some clusters being highly enriched for longer peptides. Motif plots of these submotifs, shown in Figure 3 f) (and further in Supplemental Data), exhibit submotifs that are distinct at the individual anchor residue level, and reveal that some tertiary anchors are only accessible when in conjunction with specific primary anchors, such as the P (position 3) and K (position 1, see Supplementary Figure 4) tertiary anchors not being permitted for L/A primary anchors, while the D (position 3) tertiary anchor is permitted for for all V/L/A (position 9) anchors.

**Figure 3:**
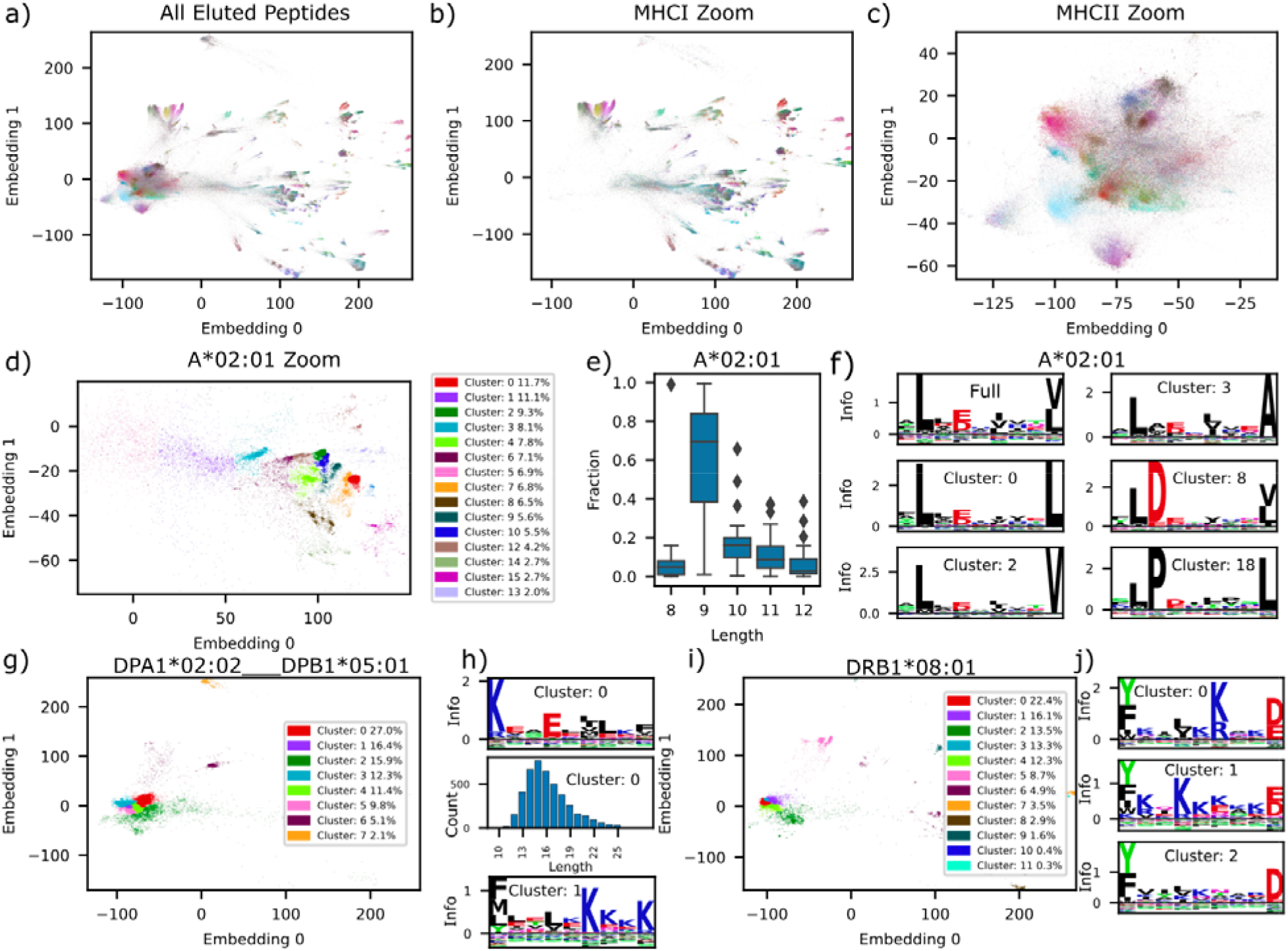
Pep2Vec’s learned peptide representation. a) Density plot of a trimap dimensional reduction of the peptide embeddings (global score 94%). The color at each pixel depicts the most common allele that is matched by the model with the peptides in that pixel. b) Depicts the data in a) but only for MHCI peptides. c) Depicts the data in a) but only for MHCI peptides. d) Density plot of the peptide embedding space zoomed into the region associated with A*02:01. Colors are generated by Louvain clustering the A*02:01 peptides in the space. Legend contains the percent of the total peptides that each cluster in d) contains. e) Boxplot depicting the fraction of each length that composes a cluster. f) Motifs depicting the overall A*02:01 motif and the motifs of various clusters depicted in d). g) Density plot of all assigned DPA1*02:02 DPB1*05:01 peptides. Clusters are generated by Louvain clustering. h) Upper: motif plot for the most populous cluster in g), middle: histogram of the peptide lengths in the upper cluster, lower: motif plot for the second most populous cluster in g) (note the inversion of the peptide). i) Density plot of all assigned DRB1*08:01 peptides. Clusters are generated by Louvain clustering. j) Three motifs depicting the various acceptable central anchor configurations for Y/F D outer anchors.

A notable achievement of Pep2Vec is its ability to generate high-quality MHCII submotifs, effectively clustering peptides based on their binding core anchors. Clear motifs are observed (see Figure 3 h,j)), suggesting that our approach of selecting the binding core starting location using the maximum calibrated BOS attention is effective, even without an explicit training reward for this behavior. Distinct clusters for inverted/non-inverted peptides, characteristic of DPA1*02:02 DPB1*05:01, are observed (Figure 3 g-h)). Inverted peptides (cluster 0) display more uniformity than non-inverted peptides (cluster 1) as indicated by the smaller spread of the cluster in the vector space. The absence of lysines at position 0 in cluster 1, a feature of inverted peptides, further differentiates these clusters. This clear separation is not achieved by other contemporary methods.^31^ These clusters also exhibit no length specificity, as shown in the histogram in Figure 3 h), because peptides of all lengths are co-located in the same cluster. Pep2Vec also uncovers subtle MHCII submotifs (Figure 3 i-j)), capturing not only the two submotifs observed by previous methods^39^ in clusters 0 and 1, but also additional submotifs, such as cluster 2.

We note briefly that manual clustering of peptides can be used to produce higher quality motifs than unsupervised clustering (such as differentiating VL anchors for submotifs with KR in position 1, and identifying subtle trends in non-anchor residues, see Supplemental Figure 4). To this end we have produced an interactive tool for exploring the Peptide latent space, enabling the user to zoom and translate through the space, and select peptides of interest for motif plotting or downstream analysis. The contaminant types found in the proceeding sections were originally identified using this tool.

### Peptide vectors can be used to detect contaminants

Through the establishment of an interpretable latent space for all peptides, regardless of allele, we can now scrutinize our MHC ligandomes with unprecedented clarity. We perform Louvain clustering over the entire latent space, which allows us to identify submotifs that are shared by multiple alleles. A significant application of this is the detection of contaminant peptides, e.g. peptides which are not actually presented by the annotated allele. These peptides pose a unique challenge, as models that better fit them achieve higher performance, despite making clearly incorrect predictions. We can identify such contaminant clusters by checking if they are dominated by a small set of samples that can potentially explain the contaminants, possibly via a different allele.

One important source of contaminant peptides is the incomplete replacement of a cell line’s original MHC allele with the new allele of interest. For example, HMy2.C1R cell lines are often used for single allelic cell line generation due to inactivating mutations in the endogenous A*02:01 and B*35:03 alleles as well as low endogenous surface expression of C*04:01.^40^ Contaminants from this residual, lowly expressed C allele are readily observable in the B*27:05 data, as shown in Figure 4 a). While most B*27:05 peptides exhibit a characteristic R/LF anchor motif (Figure 4 b)), other very distinct submotifs are also present, such as FY/D/LF motif in cluster 42 (Figure 4 c)). Upon inspection, we find that 81% of peptides from this cluster are associated with a single HMy2.C1R sample from Sabag et al.^41^, despite the existence of 26 samples for the allele. Leveraging the rest of the Peptide Vector space, we find that C*04:01 has a nearly identical FY/D/LF motif. Using the same procedure (see methods), we identify two other MHCI contaminant peptide clusters, depicted in Figure 4 d,e).

**Figure 4:**
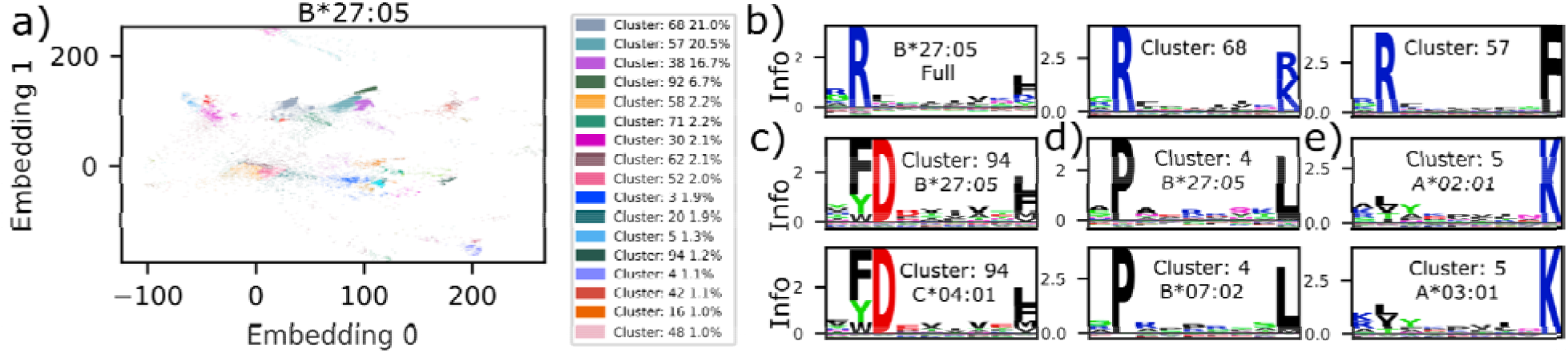
Evidence of allele contamination observed in MHCI datasets. a) Density plot of all assigned B*27:05 peptides. Clusters are generated by Louvain clustering. b) Motif plots of all B*27:05 peptides and two prominent clusters. c) Upper: Motif plot from a distinct B*27:05 cluster, and lower: a nearly identical motif from a C*04:01 cluster. d) Upper: Motif plot from a distinct B*27:05 cluster, and lower: a nearly identical motif from a B*07:02 cluster. e) Upper: Motif plot from a distinct A*02:01 cluster, and lower: a nearly identical motif from a A*03:01 cluster (see Figure 3 f) for the full A*02:01 motif).

Joint training of MHCI and MHCII allows us to identify a prevalent source of peptide contaminants, namely MHCI peptides in MHCII ligandomes. As shown earlier, the model naturally partitions the Peptide Vector space into MHCI and 2, with MHCII grouping around -75 embedding 0. When we plot only peptides from MHCII ligandomes (Figure 5 a)), we find a substantial number of peptides with embedding 0 values exceeding 0. Examining the length distribution of peptides in this region reveals that these peptides are primarily 9 amino acids long. Closer inspection of various regions reveals that the peptide motifs (Figure 5 b)) are typical of MHCI peptides, with anchor residues at positions 2 and 9, suggesting these are MHCI contaminants in MHCII ligandomes. To filter out these peptides, we trained a linear support vector machine (SVM) model on the Peptide Vectors to classify the peptide vectors for the entirety of our ligandome data as derived from either MHCI or MHCII ligandomes (see methods). After removing peptides that the SVM model classified as MHCI but were actually observed in MHCII, we noted a smoother length distribution with the removal of the peak at length 9 (Figure 5 c)). However, not all 9mer peptides were classified as MHCI, with peptides clustered around MHCII binding cores remaining unfiltered, while peptides clustered around MHCI motifs were filtered out (Figure 5 d)). Remarkably, this filtering procedure removed 6.1% of all MHCII peptides from our dataset.

**Figure 5:**
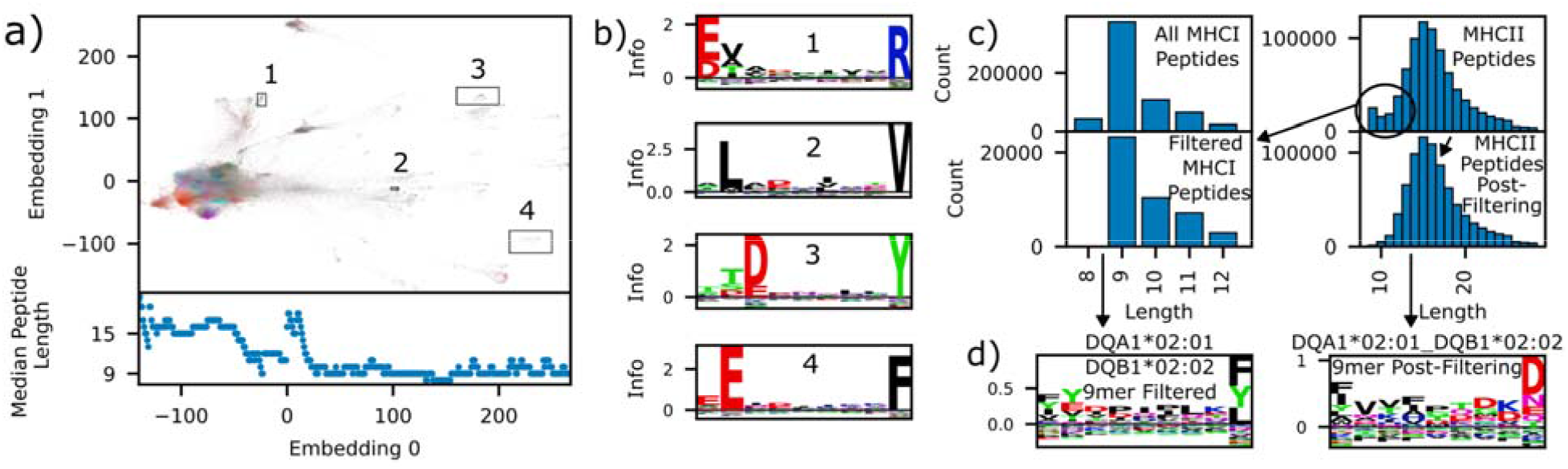
evidence of MHCI contaminant peptides in MHCII ligandomes. a) Upper: Density plot of a trimap dimensional reduction of the peptide embeddings, subset to just MHCII peptides. The color at each pixel depicts the most common allele that is matched by the model with the peptides in that pixel. Lower: the median peptide length for all peptides at a given point in the embedding 0 space. b) Motif plots from peptides contained in the four boxes drawn in a) showing anchor locations that are characteristic of MHCI. c) Upper left: histogram showing the length distribution of MHCI peptides observed in MHCI ligandomes. Upper right: histogram showing the length distribution of MHCII peptides observed in MHCII ligandomes. Lower left: histogram showing the length distribution of peptides identified as MHCI from within MHCII ligandomes by an MHCI/MHCII classifier model. Lower right: histogram showing the length distribution of peptides identified as MHCII from within MHCII ligandomes. d) Motif plot of DQA*02:01 DQB1*02:02 9mer peptides that were identified as MHCI. e) Motif plot of DQA*02:01 DQB1*02:02 9mer peptides that were identified as MHCII.

Another method for identifying contaminant peptides involves detecting unique peptides without close counterparts in the Peptide Vector space using a k-nearest neighbor (KNN) approach, shown in Figure 6 a). For each peptide, we identify its nearest neighbors and compute a motif from these peptides (or their binding cores for MHCII), identifying anchor residues and summing the KL Divergence across positions. Peptides displaying low mutual information with neighbors are considered novel. These typically reside in distinct clusters latent space, predominantly among MHCII peptides, as shown in Figure 6 b). Despite lacking a cohesive motif, these novel peptides are consistently found at protein termini (see Figure 6 c)), although protein terminal peptides are also found in the main allele clusters, indicating that the contribution from the flank embedding does not exclusively drive this behavior. Analyzing the neighbor anchor mutual information distribution reveals a significant number of MHCII peptides (and fewer MHCI peptides) exhibit very low information. We define very low information here as less than 2.5, which is motivated by the mutual information exhibited by peptides clustered only by protein termini, see Supplemental Figure 5. Examining allele DRB1*01:01 (Figure 6 e)), we find low information peptides largely do not conform to the primary motif and often originate from protein termini. We note that 80% of these peptides come from only two sources, Stražar^42^ and Marcu.^43^ The absence of a known mechanism for distinct binding of terminal peptides to MHC suggests that their overrepresentation in mass spectroscopy data is due to the simplicity of their production (only one cleavage event needs to occur to produce them), rather than an actual propensity for MHC presentation. This leads to model overfitting, as it falsely accentuates the relevance of terminal protein-derived peptides.

**Figure 6:**
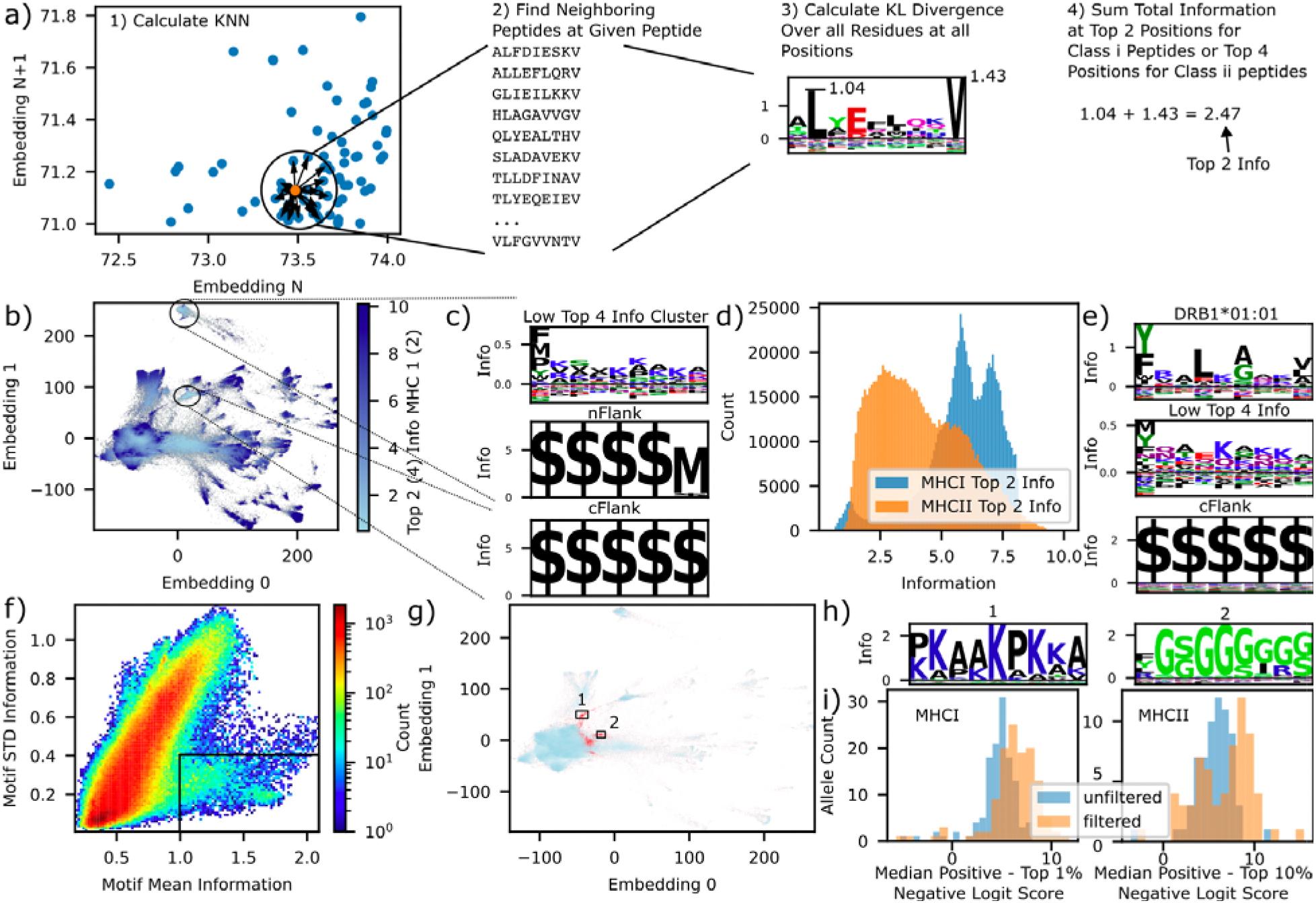
evidence of non-presented contaminant peptides in MHCII ligandomes, and Pep2Vec performance after filtering contaminant peptides. a) Schematic depicting the strategy for calculating the local mutual information of a peptide with its neighbors in the peptide embedding space. b) Density plot of the peptide space colored by the Top 2 and Top 4 position’s mutual information, for MHCI and MHCII, respectively. c) Motifs derived from peptide clusters circled in b). Upper: 9mer peptide motif from the upper cluster, middle: corresponding nFlank motif, lower: cFlank motif from the lower peptide cluster. Note the large enrichment of ‘$’ the token used to represent the end of a protein. d) Histogram of the Top 2 / Top 4 information across all peptides. e) Upper: motif plot of DRB1*01:01. Middle: motif plot of DRB1*01:01 subset to just those with less than 2 Top 4 mutual information. Lower: the corresponding cFlank to peptides in the middle plot. f) 2 d histogram of the mean and standard deviation of the motif Top 4 mutual information for MHCII. g) Density plot of MHCII peptides colored red for those in the black box in f) and light blue elsewhere. h) Motifs extracted from peptide clusters derived from those in the black boxes in g). i) Per allele histograms of the distance between the median positive logit score and the top 1% (10% for MHCII) negative logit score, showing better positive/negative separation for the model trained on filtered data, evaluated on the unfiltered test set, for both MHCI (left) and MHCII (right)

Mutual information with neighbors can also suggest contamination when excessively high. Peptides bind MHC primarily through anchor residues, while the variability in non-anchor residues is expected. This variability manifests as increased standard deviation of position-wise KL divergence values (STD) when the anchor information is high, a correlation evident in Figure 6 f). Some peptides exhibit both high mean and low STD information in their motifs, suggesting low complexity, and are grouped into three distinct clusters in the latent space, indicating a lack of allele specificity. These clusters, exclusively within MHCII, feature repetitive sequences of glycines, lysines, and alanines (Figure 6 h). Although non-specific binding to MHCII has been previously reported,^42^ these peptides markedly differ, pointing to their identity as contaminants. We note that in general many highly repetitive peptides come from common protein contaminants, such as bovine serum albumin, and are routinely filtered out of pMHC datasets, and even though we perform this filtering, the potential contaminants shown in Figure 6h were still missed.

Finally, we train a joint MHCI and MHCII peptide-presentation model (Pep2Vec Filtered) that filters out the 65,988 MHCII and 11,487 MHCI contaminant peptides identified through the interpretation strategies discussed above. The performance results are depicted in Figure 1d. A model trained on filtered data is expected to perform worse on an unfiltered test dataset because systematic contaminants can initially improve apparent model performance. To accurately evaluate the benefits of filtering, we focus on the separation between the logit scores of positive and negative samples in our test set. The lowest-scoring positive peptides from the filtered-trained model are mostly contaminant peptides present in the test set, which can skew our average precision (AP) metric. Therefore, we concentrate on the median logit scores of positive samples and compare them to the top 1% (10% for MHCII) logit scores of negative samples—these thresholds approximate the natural rate of peptide presentation by MHCI and MHCII, respectively. We find in Figure 6 i) that the median positive and top 1% negative logit score gap increases for most alleles, suggesting that the model trained on filtered data is better at distinguishing between positive and negative samples. While we find some improvement on the MHCII neoantigen dataset, in general the filtering did not improve the average precision (AP) on immunogenicity datasets, likely because AP measured ranking ability, and some highly likely presenters are not immunogenic, leading to confident models producing lower scores.

## Discussion

In developing Pep2Vec, we prioritized interpretability in order to follow recent FDA guidance for the clinical use of AI in drug discovery. As deep learning models evolve and datasets expand in size and complexity, performance gains are often achieved by capturing atypical peptides or by integrating additional features. While beneficial when these elements are reliable, experimental limitations in MHC ligandomes and biases from negative data selection can create misleading impressions of model efficacy, thus underscoring the need for interpretability. We addressed this by constructing vector spaces for all inputs that reflect our understanding of pMHC interactions. This approach enhances modularity, allowing for seamless integration of new features from protein language models (PLM).

Our analysis of protein features in Figure 2 reveals their role in determining the likelihood of peptide-MHC (pMHC) presentation. One of the dominant factors influencing a protein’s placement in the latent space is gene expression, underscoring the significant impact of peptide prevalence on MHC presentation. Although gene expression data is not explicitly provided to the model, PLM features when trained on presentation data indirectly align with expression . We also observed that learned protein representations showed further organization of the latent space by protein-protein interactions. It is not immediately clear why such a spatial organization of the protein latent space was achieved in the context of MHC presentation. Exemplary of this phenomenon: proteins involved in immune responses receive high weightings in model scores. This insight highlights a critical caveat: the protein features primarily reflect conditions from the training data studies. Significant deviations in gene expression or PPI interaction networks could diminish the utility of PLM-derived features.

Our pan-allelic, joint MHC I and II model provides the most detailed peptide map to date, surpassing previous methods like Sarkisova’s,^16^ which was limited to single peptide lengths and alleles and could not capture MHC II peptides. This comprehensive approach allows us to identify submotifs specific to certain peptide lengths (Figure 3e)) and explore how similar submotifs vary across different alleles. Particularly in our MHC II analysis (Figure 3j)), we identify a broader range of submotifs than traditional Gibbs clustering techniques.^25^ This led to our ability to identify contaminant peptides in Figure 4, which has meaningful consequences to the interpretation of overall motifs for important alleles. For example looking at the overall motif of B*27:05, one would expect that R/L is the dominant submotif, yet in reality the L anchor is artificially overweighted by contaminant submotifs, such as cluster 4.

The use of sophisticated deep neural networks in pMHC studies raises concerns, as they can inadvertently fit to non-presented peptides that appear in ligandomes due to experimental limitations, without the interpretability to recognize such errors. Previous attempts^44^ at model interpretation in pMHC, such as analyzing attention weights, have lacked the granularity needed for understanding the training data and have merely confirmed known anchor-driven presentation strategies. Our model leverages its interpretable latent space to aid detection and elimination of problematic data from the training set, preventing erroneous high-confidence predictions for similar non-presented peptides. By identifying four major sources of contamination—peptides presented by alleles assumed absent, MHC I contaminants in MHC II ligandomes, novel protein-terminus peptides, and low-complexity peptides—we have already excluded 77,475 contaminant peptides out of ∼1.5M. This list of contamination sources is not exhaustive, and future research should continue to use interpretable models to uncover additional sources.

Filtering out the contaminant peptides from MHC ligandomes used for training pMHC presentation models is essential for creating a clinic-ready model, as precious slots in personalized cancer vaccines should not be wasted on peptides that come from alleles outside a patient’s genotype, and antibodies should not be engineered to prevent the presentation of MHCI or random protein termini peptides. For this reason it is paramount to focus attention on cleaning presentation datasets, now that pMHC models are sufficiently powerful. In future work, we plan to use tools such as label smoothing^45^ to reduce the confidence of the filtered model, which led to higher scores being given to positive calls thus reducing performance on immunogenicity datasets where presented but not immunogenic peptides can occur. Additionally we plan to add new features that are predictive of immunogenicity outside of presentation to improve the filtered model in this domain.

## Methods

### Ligandome Data

All of the MHCI presented peptides used to train and evaluate models in this work are taken from our prior aggregation of publicly available MHCI ligandomes.^19^ In short, we obtained 738,427peptide:allele pairs, with 254,256 unique peptides, 347 unique genotypes and 171 unique alleles. The allele breakdown is as follows: {A: 50, B: 93, C: 28}. Only peptides from length 8-12 are used for this work.

The MHCII presented peptides used to train and evaluate models in this work were taken from two sources, our prior aggregation of publicly available MHCII ligandomes,^27^ and peptides from a recent publication that greatly expanded the available DP and DQ allotypes.^39^ In all, these datasets amount to 801,102 peptide:genotype pairs, 380,087 unique peptides, 147 unique genotypes and 83 unique alpha and beta chains. The chain breakdown is as follows: {DRA: 1, DRB1: 26, DRB3: 3, DRB4: 2, DRB5: 2, DPA1: 3, DPB1: 19, DQA1: 13, DQB1: 14}. Only peptides length 9-25 are used in this work.

These datasets are available for non-commercial use, see the data availability statement.

Mass spectroscopy of peptides is known to have a bias against measuring cysteine containing peptides. This leads to an underrepresentation of cysteines at all peptide positions, whether or not it is at an anchor residue. As a result, models trained using mass spectroscopy measured ligandomes will give peptides containing a cysteine a lower score regardless of the position it appears in. This is problematic for the model’s use in a clinical setting, where some peptides with good anchors may be given low scores because of non-anchor cysteines. In order to reduce our model’s anti-cysteine bias, we replace all cysteines in our datasets with the padding token.

### Negative Generation

Negatives used to train the model are generated via the following procedure: 1) Map all positive peptides to their source gene’s, dropping any multi mappers. 2) For each protein in the source gene, generate all possible 8-12mer peptides possible for that source gene across all proteins mapped to this source gene. 3) Drop any duplicate peptides generated from the group of proteins with a specific source gene. 4) Aggregate all peptides from all source gene’s into a single peptide table. 5) Resample this table with replacement such that every source gene has the same number of negative peptides. 6) Each epoch during training pair a random selection of peptides from this table with all alleles that exist in the positives, ensuring that negative allele distribution exactly matches that of train positives. This yields a pool of ∼46 and ∼129 million negative MHC1 and MHC2 peptides, respectively. The reference proteome used in this work was Ensembl v90.

Negatives used to test the model are generated in the same procedure, except for step 6. This last step is replaced with sampling 9x more negative peptides, for MHCII, and 99x more negative peptides for MHCI, and pairing them with the same number of alleles from test set positives, and storing them as a single table.

### Test/Train Split Development

Due to the use of protein features in Pep2Vec, we must ensure that no peptide derived from the same protein is observed in both train and test. To achieve this we follow our method developed previously,^27^ which prevents this while also achieving an even split of Gene Ontology (GO) domains and allows only minimal overlap of 9mer substring overlap between test and train. The proteins identified for this split were also used to split our MHCI peptides for this work. Splitting in this way prevents an exact test:train ratio, here, we obtained 1:7.69 for MHCI and 1:7.46 for MHCII. In our test set, we use a 1:99 positive:negative ratio for MHCI and a 1:9 positive:negative ratio for MHCII, these ratios are chosen as they are commonly used in the field to approximate the natural rate of peptide presentation, and thus give an estimate of model’s performance in a realistic setting. To eliminate any possible test set leakage, we enumerated all possible {presentation status, peptide, allele} tuples in the train and the test splits, took their intersection, and then dropped, from the test data, any {presentation status, peptide, genotype} tuples that mapped to this intersection.

### Protein Assignment, Flanking Sequences, and Gene Expression

Peptides which could not be unambiguously mapped to a single protein isoform rom our reference human proteome (GRCh38.p10) were dropped from our ligandome datasets, as in previous work.^19^ The 5 residues flanking the peptide from the source protein on both the n and c side were used as inputs into the model. In a deviation from our previous work, we did not include a special token to represent the end of a protein, and just used our padding token, as many peptides originating from the protein termini were found to be likely contaminants (Figure 6), and so inclusion of this token significantly biases the model in favor of samples with this token.

Gene expression values shown in Figure 2 d) were taken from various MHCI datasets, following our previous work.^19^ The per-protein expression values were averaged across samples.

### Immunogenicity Dataset Evaluation

MHCI cancer neoantigen datasets, Schmidt et. al.^35^ and Wells et. al.^36^ were taken as is from their publications. Flanking sequences were obtained by mapping the wildtype peptide to the human proteome (GRCh38.p10). Neoantigens that mapped to more than one source gene had all source gene-peptide-allele-flank combinations evaluated and the average score was taken as the score for the neoantigen. MHCII cancer neoantigens were taken from Reynisson et al,^37^ neoantigens were broken down into all 12-19 length peptides and predictions from these peptides were aggregated by taking the highest scoring (or lowest percentile rank) peptide for each neoantigen. Peptide:allotype combinations that were not able to be processed by MixMHC2Pred-2.2 were dropped for that model’s evaluation Clinical antibody ADA of antibody drugs were taken as is from Thrift et al.^27^ Antibody immunogenicity risk was evaluated as follows: 1) All peptides with length 12-19 are generated in a sliding window across each AB (with flanks), these peptides are paired with DRB1*01:01, DRB1*03:01, DRB1*04:01, DRB1*07:01, DRB1*08:01, DRB1*11:01, DRB1*13:01, DRB1*15:01.

Graph-pMHC is used to obtain elution likelihood scores and binding core starting predictions for each peptide-allele pair, and pairs with score less than 0 are filtered out (for NetMHCIIPan-4.3 and MixMHC2Pred-2.2, peptides rated less than weak binders are filtered out). The binding core frequency dataset is created from OASis.^46^ Graph-pMHC identified binding cores from the AB dataset are filtered out if they appear in 10 subjects. Finally, the max scoring peptide for each binding core was taken and summed across all binding cores (for MixMHC2Pred-2.2, which does not give a score, only a percentile rank, the number of binding cores is added up), and this is used as the immunogenicity risk. Antibodies are considered immunogenic if they exceed 10% population anti-drug antibody (ADA) response.

### Model Architecture and Training

All artificial neural network models in this work are implemented in pytorch,^47^ and training loops in fastai,^48^ on (which includes features such as the default fit one cycle learning rate scheduling) with mixed precision. Binary cross entropy loss and the Adam optimizer are used in their default settings. Pep2Vec uses a batch size of 3072 and a learning rate of 0.00016, except for the output multilayer perceptron (MLP), which used a learning rate of 0.00032, and the positional encoder and token embedder, which had their learning rates set to 0. Weights are initialized with default Xavier initialization. Pep2Vec is trained for 60 epochs. HLApollo’s training time on one machine with 1 NVIDIA A100 is approximately 4 hours per model. Negative set switching is implemented by sampling a new negative set as described in the negative generation section.

Pep2Vec’s Architecture is as follows. The peptide module begins with the concatenation of a dummy beginning of sequence (BOS) token. The tokens are mapped to the model dimension with a random, untrained token embedder (pytorch’s nn.embedding). A positional encoding is then added to the residues, which consists of assigning a token to each position and mapping it to the model dimension with a random, untrained token embedder (pytorch’s nn.embedding). The peptide sequence is then sent through three transformer encoder layers and the BOS representation is then extracted, and referred to as the Peptide Vector. The transformer encoder layers have custom attention masking. The first 2 layers use a sliding window mask, of size 9, in order to bias the model to predict 9mer binding cores as the start of the binding core. The last layer masks all positions, except the BOS row, since only this row is output and used by the model.

The flanking module is identical to the peptide module (with its own learned parameters), albeit with 1 transformer encoder layer instead of three. The n and c flanking residues are concatenated prior to being sent through the flanking module. This yields the Flank Vector.

The source protein features are created as follows. First, we obtained ESM-2 embeddings for all proteins from our reference human proteome using the final layer of the esm2_t36_3B_UR50D model, yielding an embedding dimension of 2560 for each residue. These embeddings were then averaged residue-wise. Then, during training, particular source protein embeddings are sent through a MLP with 1 hidden layer with the same number of nodes as the input layer, and an output layer with the same number of nodes as the model dimension. Dropout (50%) is applied after the input layer and the hidden layer. No biases are used for the final layer. This yields the Source Protein Vector.

The Peptide Vector, Flank Vector, and Source Protein Vector are added elementwise to yield the Integrated Peptide Vector.

The MHC allele features are created as follows. First, we obtained ESM-2 embeddings for all MHC in IMGT using the final layer of the esm2_t36_3B_UR50D model, yielding an embedding dimension of 2560 for each residue. These embeddings were then averaged residue-wise. Then, a principle component analysis was performed using all MHC to reduce the embedding dimension to the model dimension, creating a representation that is useful for pan-allelic generalization without using any trained features. Then, during training, particular MHC embeddings are sent through a single linear layer with the same number of nodes as the model dimension, with dropout (30%) being applied to the input embedding. This yields the MHC Vector.

Finally, the Integrated peptide Vector and the MHC Vector are multiplied elementwise and sent through a multilayer perceptron with two hidden layers. The first layer has 256 nodes and the second layer has 128 nodes. Dropout was applied after each hidden layer. The final layer is a single node that outputs the model logit score for a particular peptide:allele pair. Multiallelic deconvolution is performed using the max pair, as in our previous work.^19^

We performed a rough, manual, hyperparameter sweep over learning rate (lr), model dimension, head dropout, and transformer dropout. Presentation test set performance was used to select the hyperparameters. Batch size was fixed at 3072 to maximize the utilization of a A100 GPU, lr was tested between 0.00008 and 0.00064, 0.00016 was chosen. Model dimension was varied between {200,400, 512, 1024}, with the head dimension constant, 512 was chosen. Head and transformer dropout were varied between {0,0.1,0.2,0.4,0.5}. 0.1 and 0.3 are chosen for the head and transformer dropout, respectively. Little impact on model performance was found for most hyperparameter values (other than lr and transformer dropout). Hyperparameters not listed here were chosen based on intuition and no tuning was performed on them.

### Ensembling and Distillation

10 models were trained with different random seeds, parameter initializations, and training data ordering. These 10 models were then distilled into a single model, with double the model dimension, and consequently the latent vector dimension, to 1024. Distillation used mean squared error loss with the mean score from the 10 models for each peptide allele combination as the target score for the distilled model. The distilled model had identical evaluation metrics to the mean of the ensemble models. Distillation is performed to make inference faster, and to provide a single latent vector, without the need to perform any aggregation across individual model latent vectors.

### Binding Core Identification

Binding core’s are identified via a small set of heuristics, with the following procedure: 1) Attention maps from the last transformer stage are extracted across all attention heads, for the BOS token’s row. 2) At each position of the BOS token row the mean + 3 standard deviations of the attention values at each head is calculated and used to represent the aggregate attention across heads on a specific position. 3) This yields a vector of the same length as the peptide, finally to achieve the binding core start position, the max across the vector is used.

Optionally these binding core predictions can be calibrated in order to remove any position wise bias the model may have learned. This calibration is done by extracting the vectors yielded from step 2 for a large set of negative peptides. Then, for each peptide length a single calibration vector is extracted by averaging the attention scores across all negative samples, at each position. Using the observation that for random peptides, the binding core start position should have equal probability of being at any position, so we subtract this vector from the vector derived at step 2 above, before taking the max across positions, in step 3.

### Trimap Dimensional Reduction

Trimap^29^ was used for all dimensional reduction for plotting purposes due to its exceptional ability to maintain both local and global structure. The various hyperparameters, [n_inliers, n_outliers, n_random, weight_temp] were tuned to maximize the global score for each plot, except where the number of neighbors reaches values that are computationally infeasible. Cosine distance was always used. If multiple values with the same embedding (eg identical peptides for the peptide latent space) were in datasets, the values were uniqued prior to use in Trimap.

### Density Plots

Density plots, such as Figure 2 b-d), and Figure 3 a-d), among others, were generated using Datashader, a python library optimized for visualizing large datasets. For coloring based on categorical data, such as allele or cluster for Figure 2 a-d), the canvas is treated as a grid and the pixel color is determined by identifying the category with the largest number of data points in that pixel. For continuous data, such as in Figure 2 b-d), the average value of the data points in the pixel is calculated and the color is determined by mapping the value to the colorbar.

### Gene Ontologies

The Ensembl90 version of gene ontology terms were used in this work.

### Protein-Protein Interactions

Scanpy was used to calculate the k-nearest neighbor graph (with k=100) on the protein latent space for each protein. All protein protein interactions (PPI) with experimental evidence were obtained for each neighborhood from stringdb.^38^ The p value was calculated by producing 10,000 random neighborhoods for each protein and determining the random number of PPIs, with p = # random / # observed, making a lower limit estimation of 10^-5^.

### Clustering

Leidan clustering was used for all clustering in this work. Scanpy was used to calculate the k-nearest neighbor graph (with k=100), and the leidenalg library was used to calculate the partitions using RBConfigurationVertexPartition. Submotif cluster assignment was done on all the data for just one allele at a time, with multiallelic data being assigned to a particular allele by the model. This was performed using all data, train and test. A resolution parameter of 0.1 was used for MHCI alleles, and 0.3 was used for MHCII alleles. For Figure 4 a where the entire latent space was clustered together, a resolution parameter of 0.1 was used. For the low complexity peptides in Figure 6 h) k=20 was used, and the resolution parameter was also set to 0.1. These parameters were chosen by eye.

### TM-Align Score

TM align scores were calculated using foldseek^49^ on structures from AlphfoldDB^50,51^. Neighborhood scores were calculated using the 100 nearest neighbors from the Protein latent k-nearest neighbor latent space.

### Zoom Plots

Figures 3b-d) do not depict all data corresponding to their condition (MHCI/MHCII/A*02:01), and instead limit to just the embedding 0 and 1 coordinates depicted in the plot, and are hence called zoom plots. This is done to avoid showing contaminant data points which are shown later in Figures 4 and 5, which improves the readability of the figure, and the flow of the manuscript. Data subsets in Figure 5 b) are chosen by eye based on dense and diverse regions found in the MHCI contamination space. The data subset in Figure 6 c) is chosen by eye as representative of the low mutual information region.

### Motif Logos

Motif logos are plotted using the logomaker library.^52^ The KL Divergences are calculated relative to the baseline probabilities of each amino acid as found in our reference human proteome (Ensembl v90).

### Peptide Latent Space Visualization Tool

We provide a visualization tool for navigating the large space of peptide vectors extracted from the model for the research community to find other potential contaminants or biases. This tool is created using the holoviews python tool set and library. Specifically the datashader library is used to rasterize, color, and plot the reduced latent space. Panel is used for the user interface controls and layout. Finally, motif plots for selected peptides are provided using the logomaker library.^52^ Source code and usage instructions are provided in the public github for this project.

### Allele Contamination Identification

Allele contamination as found in Figure 4 as follows. For each cluster in the peptide latent space (clusters were assigned using all alleles) each allele presenting peptides in the cluster was considered. For each allele the number of presented peptides divided by the total number of presented peptides obtained for that allele (the normalized peptide count) was identified, as well as the number of samples producing peptides within that cluster divided by the total number of samples obtained for that allele (the normalized dataset count). Then, the allele with the most peptides in that cluster (the max allele) was identified, and the similarity between the allele and the max allele was obtained by cosine similarity of the MHC representation used by the model. The max alleles normalized peptide count was also identified. Allele:cluster tuples with a normalized peptide count <10%, a sample fraction <33%, a max allele normalized peptide count >35%, a similarity of <90%, and at least 100 peptides within the cluster were flagged as likely contaminants. This produced 22 flagged allele:clusters from 14 clusters.

### MHCI/II Contamination Identification

MHCI/II assignment was performed by training a linear support vector machine (SVM) classifier (implemented in sklearn) that takes in Peptide Vectors and classifies them as either MHCI or 2. A 10 fold, kfolds strategy was used to get predictions from all data points in order to make sure predictions are always made on held out examples from perspective of the classifier. Peptides that were assigned to the wrong MHC type were considered contaminants.

### Nearest Neighbor Mutual Information Calculation

We use a proxy metric derived from kNN mutual information in order to assess the accuracy of binding core prediction, as well as the models ability to place similar peptides near each other in the latent space. We calculate this information content metric for MHCII examples via the following procedure: 1) For each positive peptide, find its nearest 1000 neighbors in the model full vector latent space, using cosine distance metric. 2) We then extract the 1000 predicted binding core’s, then, using logomaker, extract the number of bits of information per position by summing the number of bits at each position for each amino acid, across amino acids. 3) Then the highest 4 positions are summed together to create a single score.

We calculate this metric for MHCI in a similar procedure, except the procedure is run for each peptide length independently, and the top 2 positions are summed, instead of 4.

### Information-based Contamination Identification

Low mutual information peptides were considered contaminants if their top 2 or top 4 information scores were less than 1.25.

Low complexity peptides were considered contaminants if they had a position-mean KL divergence of at least 1.25, and a position-wise standard deviation of less than 0.5. These thresholds were chosen by eye using the plot if Figure 6 f).

### Large Language Model Usage

ChatGPT (version 4 and 4o) was used to edit the language of this manuscript for clarity, concision, and flow. ChatGPT was also used to identify areas of the text where inconsistent language was used, redundancy occurred, and where key findings were not adequately highlighted so that they could be improved. ChatGPT was always given already written text to improve, and was not asked to generate its own ideas. All text generated by ChatGPT was carefully edited for factual accuracy.

## Data Availability Statement

All datasets utilized in this study, including those used for benchmarking are comprehensively provided and readily accessible at: https://zenodo.org/records/13930613. Researchers and interested parties are encouraged to explore and utilize these datasets in accordance with the guidelines provided in the repository.

## Code Availability Statement

Downloading, installing, or using Pep2Vec is subject to agreement to the terms of the Pep2Vec License attached herewith. The executable is available at https://github.com/Genentech/Pep2Vec. This github also contains Pep2Vec_viz an interactive visualization tool for the peptide latent space created by Pep2Vec. Please note that the Pep2Vec licenses allows Pep2Vec to be downloaded, installed, and/or used for internal teaching and non-commercial academic research purposes only.

---

**Supplementary Figure 1:**
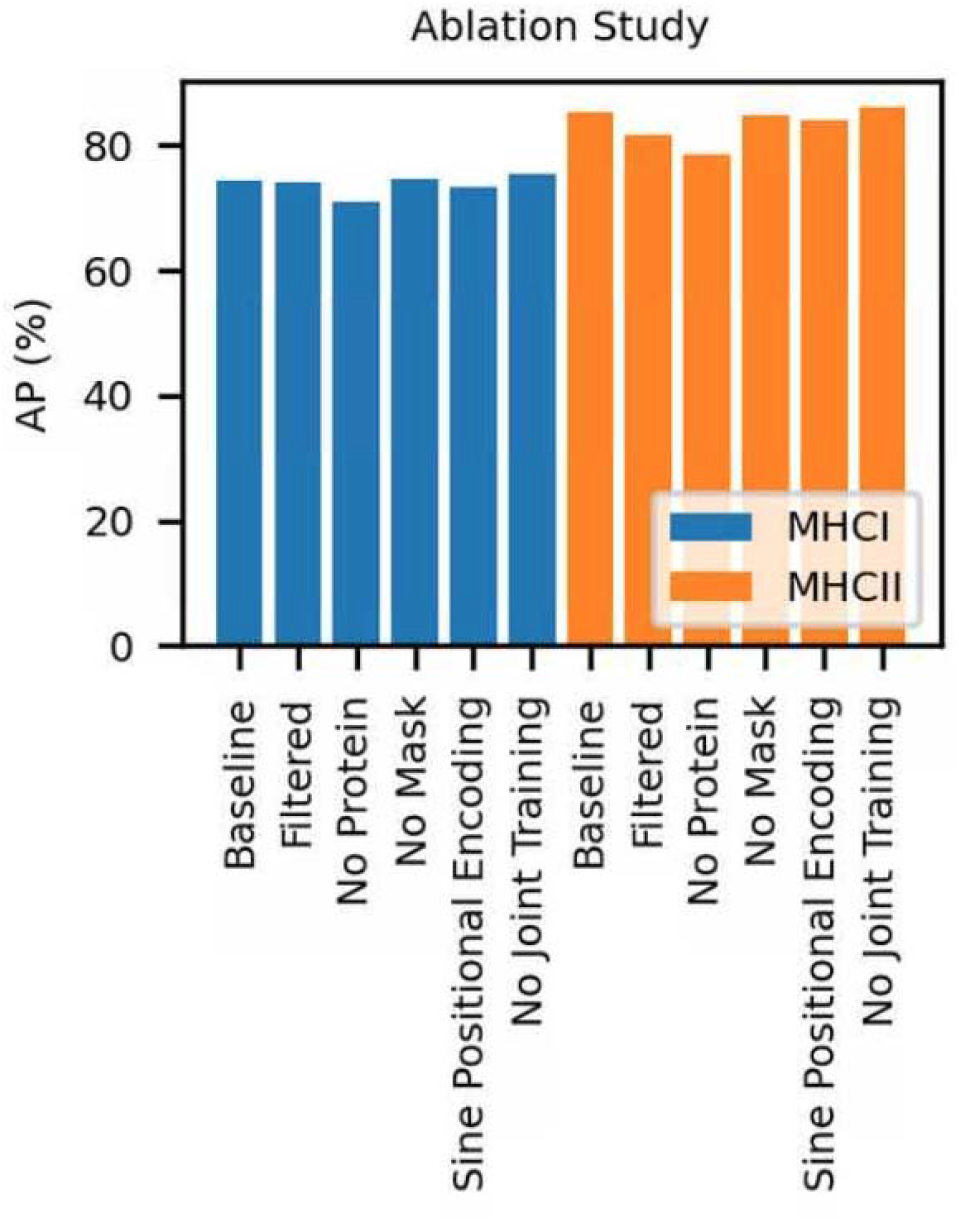
ablation study of features and training techniques added in this work.

**Supplementary Figure 2:**
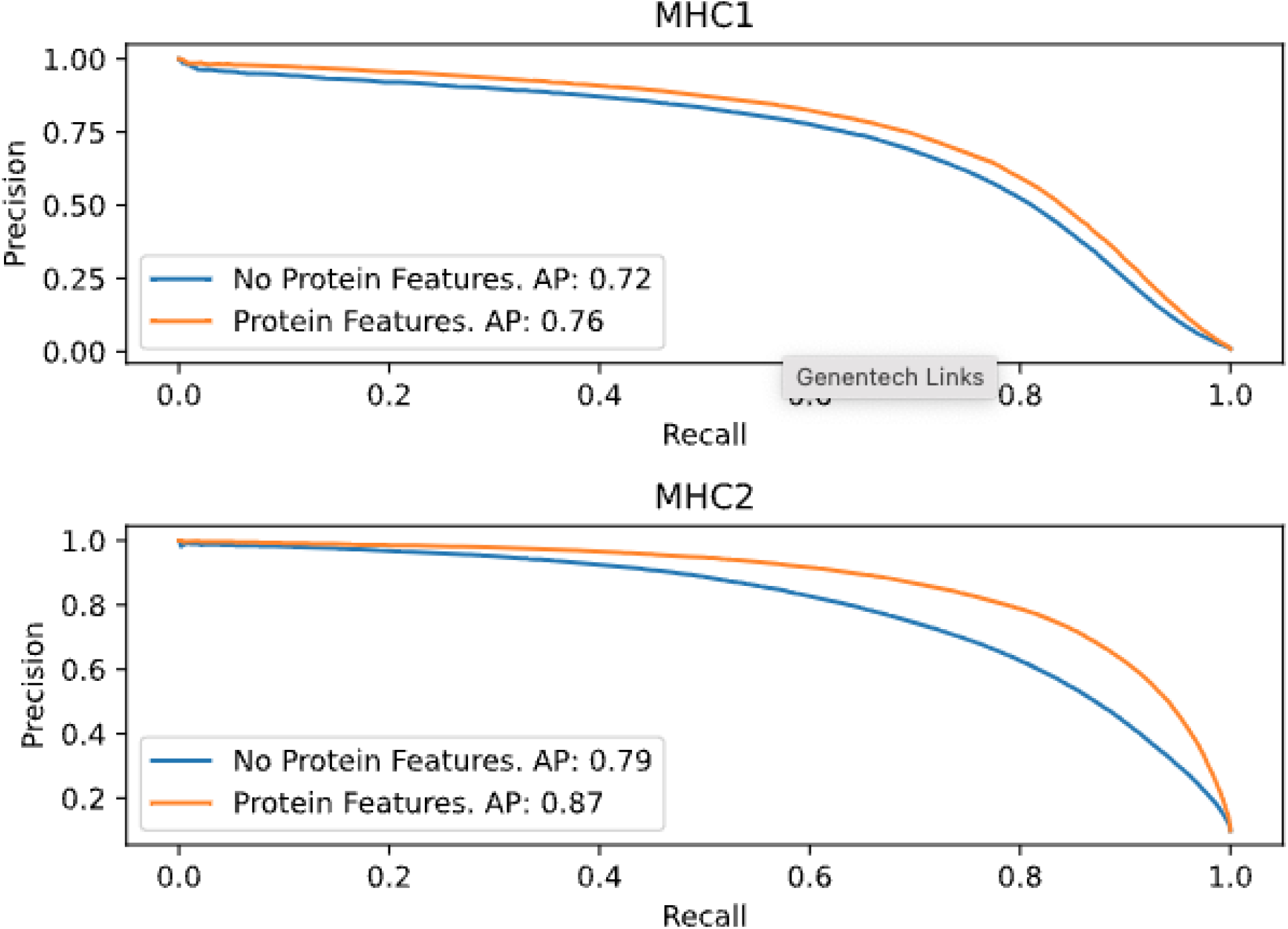
Precision-recall curves for models trained with and without protein features. Protein features add meaningful improvements to precision over all recall values.

**Supplementary Figure 3:**
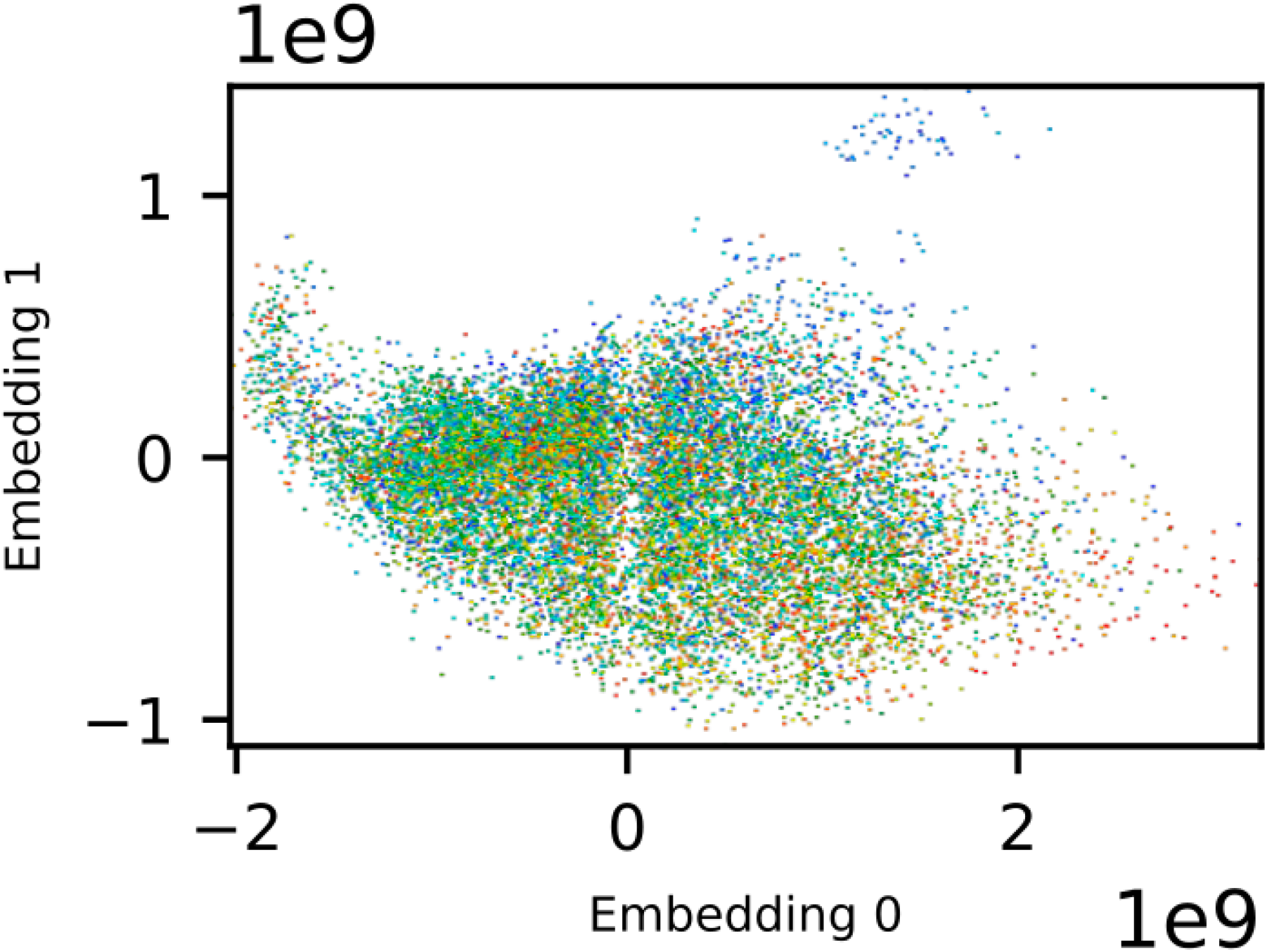
A density plot of a trimap dimensional reduction (global score 98%, indicating good dimensional reduction with respect to the original source features) of the raw ESM embeddings for all source proteins in our dataset. The color depicts the log transformed expression value of each protein, averaged by the proteins at each particular pixel of the plot. Expression values are themselves an average of the expression values observed across various MHCI datasets.

**Supplementary Figure 4:**
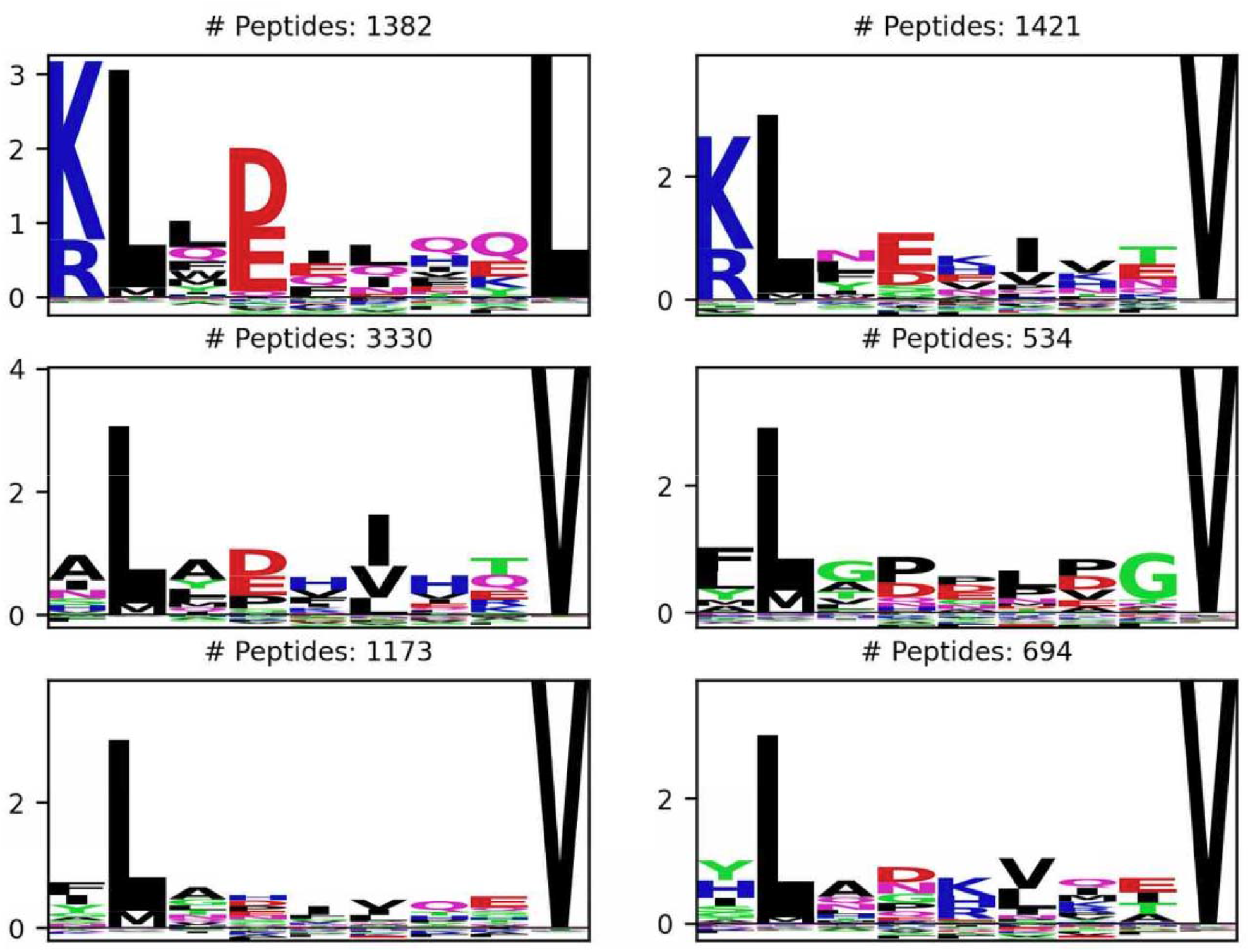
Submotifs of A*02:01 manually identified using our visualization tool, pep2vec viz. Extremely sensitive detection of patterns outside the primary anchors are observed.

**Supplementary Figure 5:**
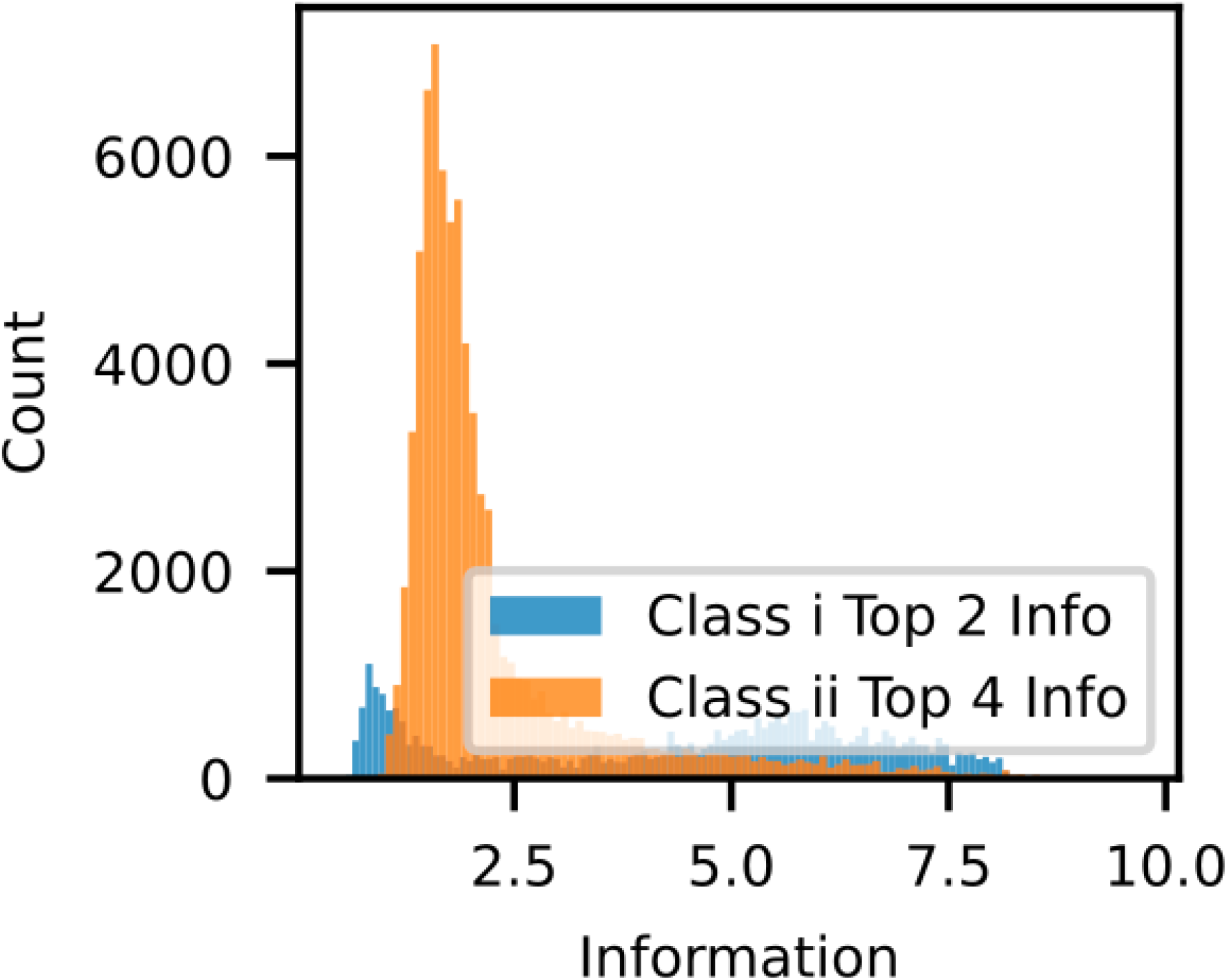
Histogram of the Top 2 / Top 4 information across protein termini peptides. Note that most have mutual information under about 2.5.

## Notes

https://zenodo.org/records/13930613

https://github.com/Genentech/Pep2Vec

### Competing Interest Statement

All authors are employees of Genentech

